# Perception of whole-body motion during balance perturbations is impaired in Parkinson’s disease and is associated with balance impairment

**DOI:** 10.1101/666636

**Authors:** Sistania M. Bong, J. Lucas McKay, Stewart A. Factor, Lena H. Ting

## Abstract

**Background:** In addition to motor deficits, Parkinson’s disease (PD) may cause perceptual impairments. The role of perceptual impairments in sensorimotor function is unclear, and has typically been studied in single-joint motions. Research Question: We hypothesized that perception of whole-body motion is impaired in PD and contributes to balance impairments. We tested 1) whether directional acuity to whole body perturbations during standing was worse in people with PD compared to neurotypical older adults (NOA), and 2) whether balance ability, as assessed by the MiniBESTest, was associated with poor directional acuity in either group.

**Methods:** Participants were exposed to pairs of support-surface translation perturbations in a two-alternative forced choice testing paradigm developed previously in a young healthy population. The first perturbation of each pair was directly backward and the second deviated to the left or right (1°–44°). Participants judged and reported whether the perturbations in each pair were in the “same” or “different” direction. This information was used to calculate directional acuity thresholds corresponding to “just-noticeable differences” in perturbation direction. Linear mixed models determined associations between directional thresholds and clinical variables including MDS UPDRS-III score, age, and MiniBESTest score. Results: 20 PD (64±7 y, 12 male, ⩾12 hours since last intake of antiparkinsonian medications) and 12 NOA (64±8, 6 male) were assessed. Directional thresholds were higher (worse) among PD participants (17.6±5.9° vs. 12.8±3.3°, P<0.01). Linear mixed models further showed that higher thresholds were associated with MDS UPDRS-III score (P<0.01), and were associated with poorer balance ability among PD participants (P<0.01), but not among NOA participants (P=0.40). Significance: Perception of whole-body motion is impaired in PD and may contribute to impaired balance and falls.

## Introduction

Deficits in somatosensory perception have been identified in Parkinson’s disease (PD) [1-8], but their relationship to impaired balance is unclear. PD causes profound balance problems and falls [9], resulting in significant morbidity, mortality, healthcare expense, and reduced quality of life [10-12]. Because standing balance requires complex sensorimotor integration [13-15], impairments in the ability to accurately sense body motion would be expected to contribute to balance impairments. In laboratory studies using perturbation testing, balance-correcting muscle responses evoked by perturbations are less directionally-specific in people with PD than in matched individuals [16, 17], which could be explained by an impairment in the ability to accurately sense the direction of body motion. Our goal was to explicitly test whether perception of whole-body motion direction is impaired in individuals with PD, and if so, to test whether this impairment is related to impaired balance.

Somatosensory perceptual deficits in PD have been typically investigated in single-joint tasks [3, 7, 18-20] usually of isolated limbs [21-23]. Results demonstrate impaired perception of the position of single joints including the ankle [20], elbow [6, 7], and index finger [25] as well as of the rotation of the trunk [24]. Far fewer studies have examined perceptual deficits in PD during walking and standing. When asked to estimate the height of obstacles encountered during walking using only proprioceptive information from the feet and legs, people with PD overestimate obstacle height, suggesting that proprioceptive feedback may be impaired [25].

Additionally, in people with PD and freezing of gait, perception of gait speed differences between the legs while walking on a split-belt treadmill is less accurate compared to those without freezing [26]. However, whether individuals with PD have impaired perception of whole-body motion direction during standing remains unknown.

We previously developed a balance testing paradigm to quantify the perception of whole-body motion direction during backward perturbations to standing balance in young healthy subjects using a “Parameter Estimation by Sequential Testing” (PEST) algorithm [27, 28]. We quantified just-noticeable differences to lateral deviations during backward support-surface translations while standing using a two-alternative forced choice paradigm, and identified a directional threshold of about 10° [28]. We demonstrated that the PEST algorithm required far fewer trials than conventional psychophysical methods, making the testing paradigm feasible in clinical populations like PD.

In this study, we quantified directional acuity of whole-body motion during backward perturbations to standing balance in individuals with PD, and compared it to that of matched neurotypical older adults. We hypothesized that 1) whole-body motion directional acuity would be impaired in people with PD, and 2) impaired whole-body motion directional acuity would be associated with impaired balance, as assessed with a behavioral clinical balance scale in common clinical use in PD, the MiniBESTest [29, 30].

## Methods

### Participants and setting

We performed a cross-sectional study of whole-body motion perception in community-dwelling individuals with PD and age- and sex-similar neurotypical older adults (NOA). Participants were recruited from the Jean & Paul Amos Parkinson’s Disease and Movement Disorders Clinic at Emory University School of Medicine, community outreach events, senior centers and similar venues. PD subjects were recruited from the study population of an observational 1-year fall risk study. NOA participants were recruited for a one-time visit. Sample size for both groups was selected to meet or exceed general guidelines of N=12 recommended for pilot studies [31].

All participants were age ≥40 years without a history of musculoskeletal and/or neurological disorders, other than PD, sensory deficits, or dizziness. Participants with PD were diagnosed with idiopathic PD and cleared for participation in the study by their neurologist. All participants provided informed consent prior to participation according to a protocol approved by Institutional Review Board of Emory University.

### Study visit

Informed consent and all testing procedures were performed during a single ≈3 hour study visit. PD participants were assessed in the practically defined OFF state [32], which is ≥12 hours after last intake of antiparkinsonian medications. Demographic and clinical information were collected using standardized instruments. These included Movement Disorders Society-Unified Parkinson’s Disease Rating Scale (MDS-UPDRS) Parts I-IV [33], MiniBESTest [34]for balance ability, Montreal Cognitive Assessment (MoCA) [35]) for cognition, and Freezing of Gait Questionnaire (FOG-Q) [36]. The motor portion of the MDS-UPDRS (MDS-UPDRS-III) was scored from video by a movement disorders specialist (S.A.F.). PD disease duration and other clinical information were obtained from clinical records when possible and otherwise by interview. The number of falls over the preceding 6 months were quantified by self-report. Antiparkinsonian medications were summarized as levodopa equivalent daily dose (LEDD) according to standardized formulae [37].

### Whole-body motion perception testing

Participants stood barefoot on a custom perturbation platform that translated in the horizontal plane, wore a harness attached to the ceiling, and were attended by staff [17, 38]. Stance width between the feet was standardized to inter-anterior superior iliac spine (ASIS) distance. Participants were blindfolded and wore headphones that played white noise to eliminate visual and auditory feedback during testing. All testing was performed while participants were instrumented with electromyography electrodes on the legs and reflective motion capture markers [17, 38].

Directional acuity threshold of a whole-body motion was defined as the smallest detectable difference of lateral deviations between pairs of sequential perturbations during standing (Figure 1). Each trial consisted of a pair of two ramp-and-hold translations of the support surface (7.5 cm displacement,15 cm/s peak velocity, 0.1 m/s^2^ peak acceleration) [17, 38]. The first perturbation of each pair was delivered in the backward direction, causing the body center of mass to be displaced anteriorly with respect to the ankles. The second perturbation of each pair was delivered 0.5 s after the first perturbation ended, and deviated in a pseudo-random fashion either to the left or to the right from the backward direction, and by an amount Δ*θ* determined adaptively online using the PEST with identical parameters as Puntkattalee et al [28], with an initial Δ*θ* of +3° or -3°, an initial step size of 4°, and a stopping step size criterion of 0.5°.

**Figure 1.**
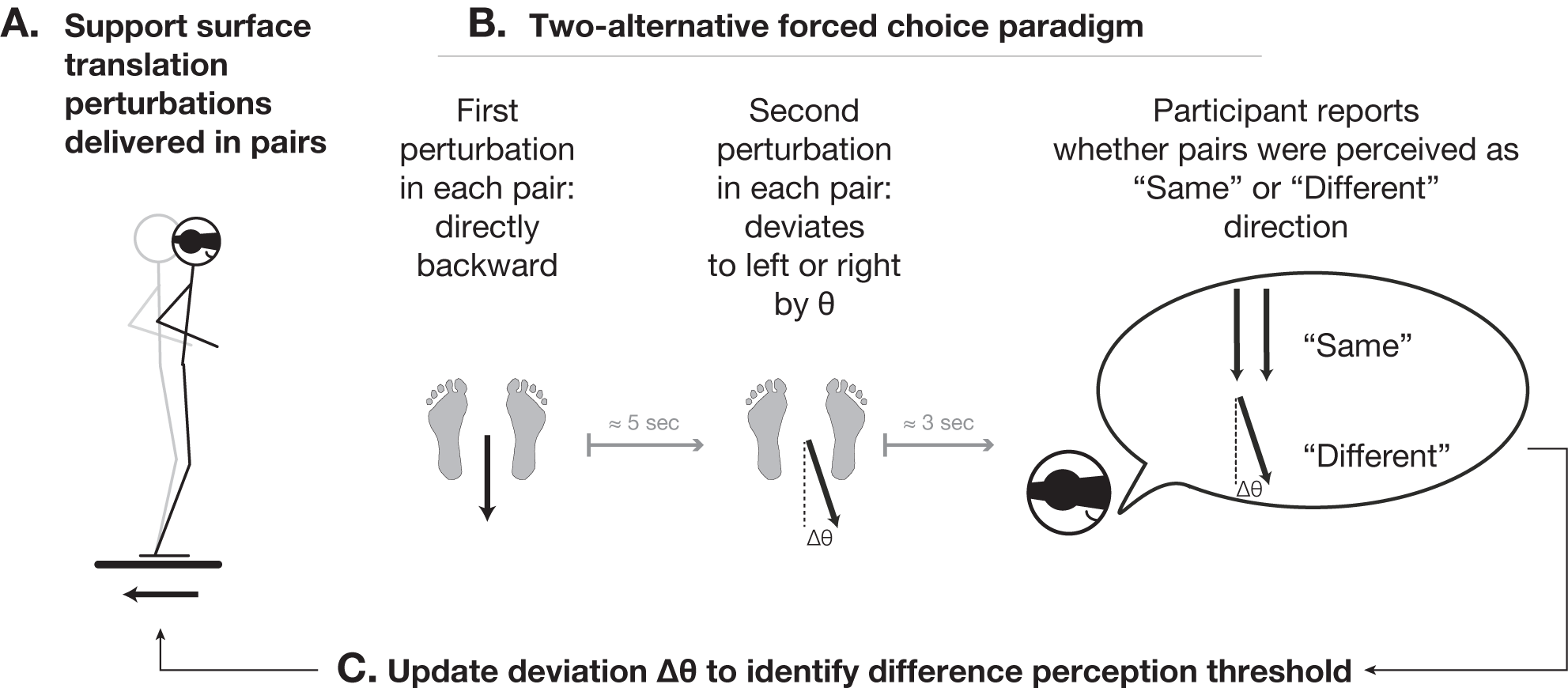
Schematic illustration of experimental paradigm and Parameter Estimation by Sequential Testing (PEST) algorithm. A. Perturbations were delivered as in earlier studies in individuals with PD [17, 38]. B. Participants judged and reported whether both perturbations delivered in pairs that were identical in magnitude but that were separated by a deviation angle Δ*θ* were in the “Same” or “Different” direction. C. Between each pair of perturbations, the deviation angle Δ*θ* was altered adaptively in order to identify the just noticeable difference threshold Δ*θ* _*Threshold*_.

Shortly after each pair of perturbations (<3 seconds), participants judged and reported verbally whether the two perturbations in the pair were in the “same” or “different” direction. Verbal responses from patients with hypophonia were repeated by a nearby study staff member for recording as necessary. During testing, participants held a small push-button device to enable comparisons with earlier results[28]; signals from the handheld device were not examined.

At the beginning of each testing session, participants were given 4-5 practice trials to ensure they understood the task. Trials in which participants stepped in response to the perturbation were excluded from analysis. Convergence to threshold for right and left deviations were found simultaneously by interleaving left and right perturbations during data collection. Up to 60 pairs of perturbations were administered to each participant, with data collection ending earlier when the iterative step size was reduced to ⩽0.5°.

## Data analysis

Threshold values Δ*θ*_*Threshold*_ were obtained from the left and right side of each participant and were reclassified in ascending order of magnitude as minimum and maximum. In cases where convergence was not reached, threshold values were estimated using a psychometric curve fit to the PEST response data (Supplementary Materials S1).

## Statistical analysis

Differences in demographic and clinical features between PD and NOA groups were assessed with chi-squared tests and independent samples *t*-tests as appropriate. Additional *t*-tests evaluated perceptual asymmetry between maximum and minimum thresholds in each participant and across groups. Differences between maximum and minimum threshold values were log-transformed prior to analysis in order to meet criteria for approximate normality as determined by the Kolmogorov-Smirnov test.

Overall differences between PD and NOA in identified threshold values were assessed using independent samples *t*-tests. Multivariate linear mixed models were used to further examine the contributions of age and parkinsonian severity to the perception of whole-body motion. We fit the following model to each of the maximum and minimum thresholds Δ*θ*_*Threshold*_:

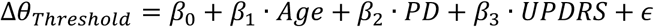

Where Δ*θ* _*Threshold*_ corresponds to the identified threshold value from either side of the body, *Age* corresponds to age in years, *PD* is a dichotomous variable coding the presence of PD, and *UPDRS* specifically represents MDS-UPDRS-III (motor examination) score. MDS-UPDRS-III scores were centered about zero prior to entry in linear mixed models.

To test hypothesis 1: whether whole-body motion directional acuity is impaired in people with PD, the null hypothesis *β*_2_ = 0 was evaluated with an F-test.

To test hypothesis 2: whether impaired perception of whole-body motion direction is associated with impaired balance deficits, we constructed linear mixed models that regressed observed MiniBESTest scores onto observed threshold values, with interaction terms that allowed for different regression coefficients among the NOA and PD groups. Separate models were constructed for each of the maximum and minimum thresholds, structured as follows:

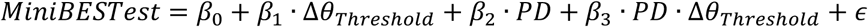

To test whether impaired perception of whole-body motion direction was associated with balance deficits, we evaluated the following null hypothesis using F-tests. First, to test whether impaired perception was associated with balance deficits among NOA, the reference group, we evaluated *β*_1_; = 0. Second, to test whether impaired perception was associated with balance deficits among PD, we evaluated *β*_1_;+ *β*_3_= 0. Finally, to test whether associations were significantly different among the NOA and PD groups, we evaluated *β*_*3*_; = 0.

## Additional analyses

Additional analyses were performed, including analyses of: 1) Differences in threshold values between PD/NOA participants and existing data of young healthy adults [28], 2) Clinical and demographic features associated with failure to complete testing protocol, and, 3) Clinical and demographic features associated with incomplete threshold convergence during the testing protocol, 4) Associations between MDS-UPDRS-III motor symptom asymmetry [39] and threshold values, and, 5) Whether direction threshold magnitude influenced the likelihood of convergence (Supplementary Materials S1).

## Results

### Demographic and clinical characteristics

Demographic and clinical characteristics of PD (N=20, 12 male, age 64±7 y, disease duration 8±5y) and NOA groups (N=12, 6 male, age 64 ± 78 y) are summarized in Table 1. No significant group differences were identified between PD and NOA in sex, age, height, weight, or MoCA score. Age ranges were 56–80 (PD) and 56–81 (NOA). PD patients had significantly poorer performance on MiniBESTest (3.5 points, p=0.0018). Average MoCA scores were indicative of normal cognition (>27) in both groups. 19/20 PD participants were prescribed antiparkinsonian medications at the time of enrollment; 1 participant had not yet begun pharmacotherapy.

**Table 1.**
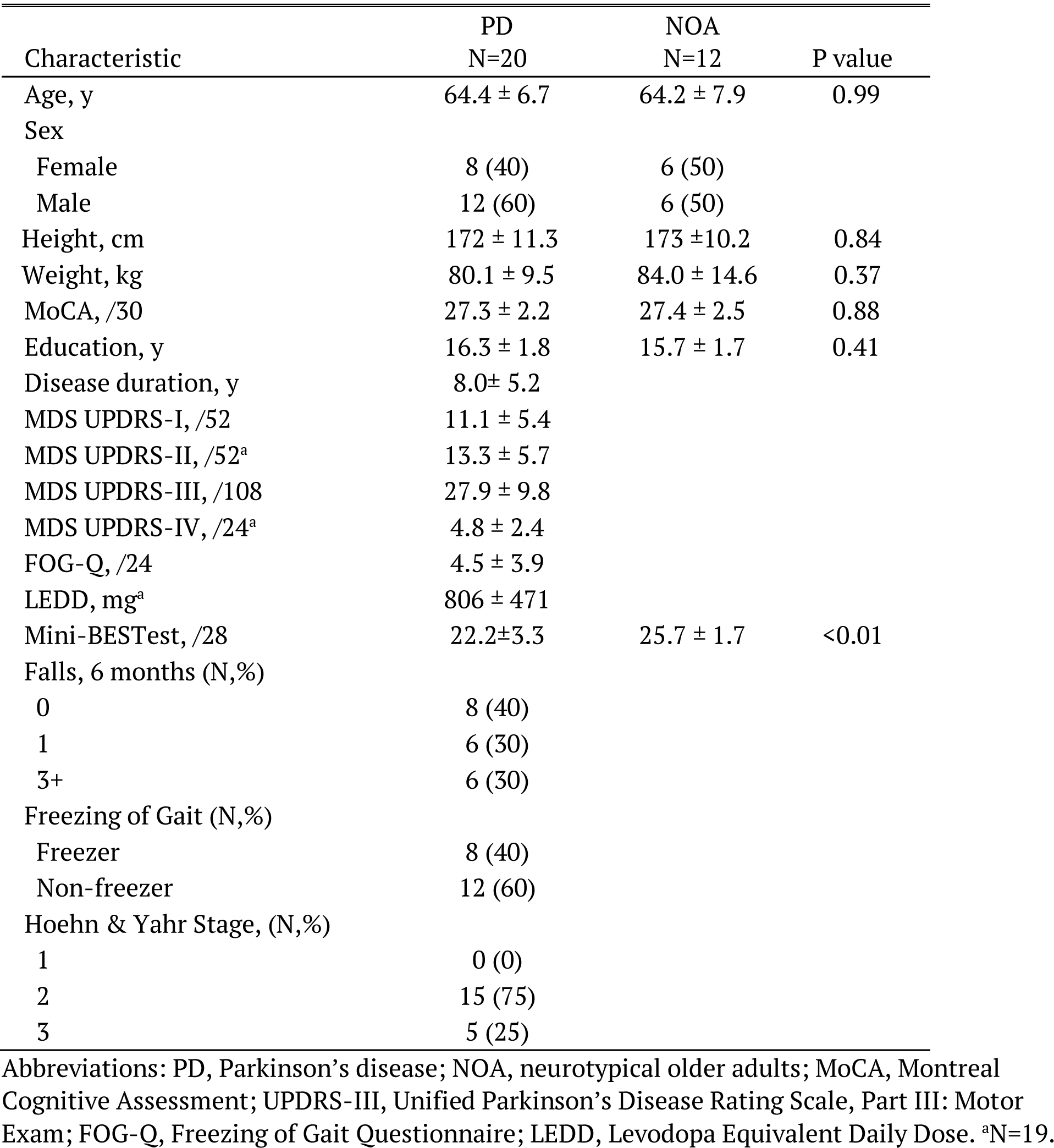
Demographic and clinical characteristics of study participants.

| Characteristic | PD<br>N=20 | NOA<br>N=12 | P value |
| --- | --- | --- | --- |
| Age, y | 64.4 ± 6.7 | 64.2 ± 7.9 | 0.99 |
| Sex |  |  |  |
| Female | 8 (40) | 6 (50) |  |
| Male | 12 (60) | 6 (50) |  |
| Height, cm | 172 ± 11.3 | 173 ± 10.2 | 0.84 |
| Weight, kg | 80.1 ± 9.5 | 84.0 ± 14.6 | 0.37 |
| MoCA, /30 | 27.3 ± 2.2 | 27.4 ± 2.5 | 0.88 |
| Education, y | 16.3 ± 1.8 | 15.7 ± 1.7 | 0.41 |
| Disease duration, y | 8.0 ± 5.2 |  |  |
| MDS UPDRS-I, /52 | 11.1 ± 5.4 |  |  |
| MDS UPDRS-II, /52<sup>a</sup> | 13.3 ± 5.7 |  |  |
| MDS UPDRS-III, /108 | 27.9 ± 9.8 |  |  |
| MDS UPDRS-IV, /24<sup>a</sup> | 4.8 ± 2.4 |  |  |
| FOG-Q, /24 | 4.5 ± 3.9 |  |  |
| LEDD, mg<sup>a</sup> | 806 ± 471 |  |  |
| Mini-BESTest, /28 | 22.2 ± 3.3 | 25.7 ± 1.7 | <0.01 |
| Falls, 6 months (N,%) |  |  |  |
| 0 | 8 (40) |  |  |
| 1 | 6 (30) |  |  |
| 3+ | 6 (30) |  |  |
| Freezing of Gait (N,%) |  |  |  |
| Freezer | 8 (40) |  |  |
| Non-freezer | 12 (60) |  |  |
| Hoehn & Yahr Stage, (N,%) |  |  |  |
| 1 | 0 (0) |  |  |
| 2 | 15 (75) |  |  |
| 3 | 5 (25) |  |  |

Abbreviations: PD, Parkinson's disease; NOA, neurotypical older adults; MoCA, Montreal Cognitive Assessment; UPDRS-III, Unified Parkinson's Disease Rating Scale, Part III: Motor Exam; FOG-Q, Freezing of Gait Questionnaire; LEDD, Levodopa Equivalent Daily Dose. <sup>a</sup>N=19.

In addition to these participants, testing was attempted but not completed in N=14 additional participants (N=10 PD, N=4 NOA), due to: fatigue (N=4 PD); inability to tolerate headphones and blindfold (N=4 PD, N=1 NOA); inability to understand instructions (N=1 PD, N=2 NOA), and testing equipment malfunction (N=1 PD, N=1 NOA). Neither the frequency of nor the reasons for early termination varied across groups (P=0.50 and P=0.28, respectively). Inability to complete testing was associated with increased age (p=0.02) and impaired MiniBESTest performance (p=0.017) but not with presence of PD (p=0.44) or other variables related to PD severity (Supplementary Materials S1).

### Whole-body directional acuity is impaired in PD

Independent samples *t*-tests showed that directional acuity thresholds were significantly larger in PD vs. NOA (maximum Δ*θ*_*Threshold*_, 17.6±5.9° vs. 12.8±3.3°, p=0.016; minimum Δ*θ*_*Threshold*_, 13.5±4.0° vs. 9.67±2.9°, p=0.008; Figure 2A,B). Differences between maximum and minimum Δ*θ*_*Threshold*_ were statistically-significant for both groups (p<0.001). No group differences in this perceptual asymmetry were identified (p=0.66).

**Figure 2.**
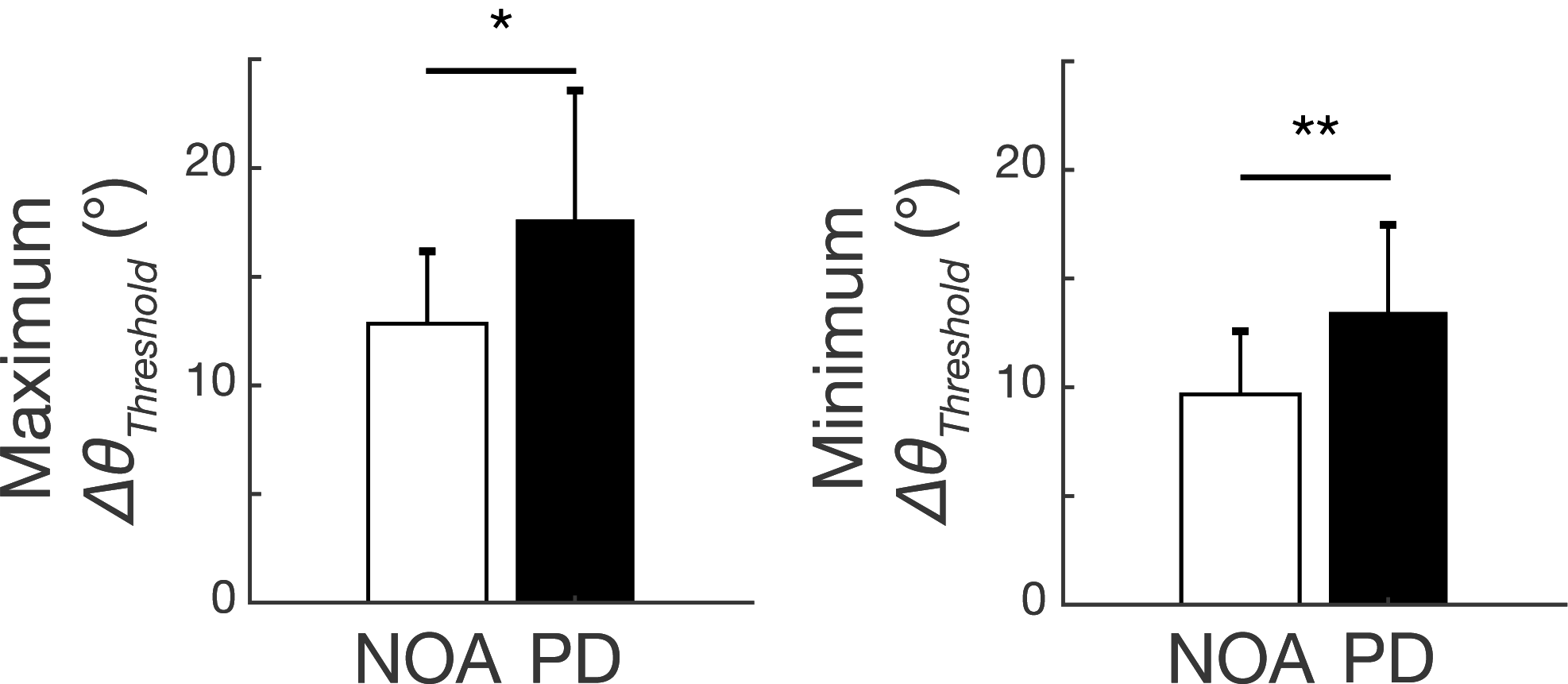
Comparison of whole-body motion direction perception in NOA and PD. Left: Maximum Threshold. Right: Minimum Threshold. *P<0.05, **P<0.01, *t*-test.

Linear mixed models showed that the presence of PD, and PD severity as measured by MDS-UPDRS III, were both significantly associated with impaired perception of whole-body motion (maximum Δ*θ*_*Threshold*_: PD presence, p<0.001, PD severity, p<0.01; minimum Δ*θ* _*Threshold*_: PD presence, p<0.01, PD severity, p<0.01). No significant effects of age were identified (maximum Δ*θ*_*Threshold*_: p=0.32, minimum Δ*θ* _*Threshold*_: p=0.82) (Table 2).

**Table 2.** Associations between whole-body motion perception, PD, and age determined by linear mixed models.

| Predictor | Beta coefficient | 95% CI | P value |
| --- | --- | --- | --- |
| Maximum Threshold |  |  |  |
| Age (°/y) | -0.11 | -0.34, +0.11 | 0.315 |
| PD (°) | +4.81 | +1.71, +7.91 | 0.004** |
| MDS-UPDRS-III (°/point) | +0.43 | +0.22, +0.63 | <0.001** |
| Minimum Threshold |  |  |  |
| Age (°/y) | -0.02 | -0.20, +0.17 | 0.822 |
| PD (°) | +3.78 | +1.21, +6.36 | 0.006** |
| MDS-UPDRS-III (°/point) | +0.21 | +0.03, +0.39 | 0.045* |

\*P<0.05, \*\*P<0.01, significant effect of predictor variable on threshold value, linear mixed models.

In ten participants who completed the planned testing protocol (N=7, PD; N=3, NOA), the algorithm failed to “converge” – to settle upon a final measurement for threshold value – on one side during the allotted time. These threshold values were estimated using a psychometric curve fit to the PEST response data (Table S1). The frequency of convergence failure did not vary across groups (p=0.71). No statistically-significant associations were identified between convergence failure and any clinical or demographic variables examined. The strongest potential association was with impaired MiniBESTest performance (p=0.22; Supplementary Materials S1).

### Worse balance is associated with impaired directional acuity among PD but not NOA participants

Linear mixed models examining associations between MiniBESTest score and directional acuity identified statistically-significant group×threshold interaction terms for both maximum (p=0.017) and minimum (p=0.007) Δ*θ* _*Threshold*_ (Figure 3A,B). Among PD participants, associations between MiniBESTest score and directional threshold were -0.40 points/° (p<0.001) and -0.53 points/° (p=0.002), respectively (Table S2). No statistically-significant associations between directional perception and MiniBESTest score were identified among NOA (p>0.43; Supplementary Materials S1).

**Figure 3.**
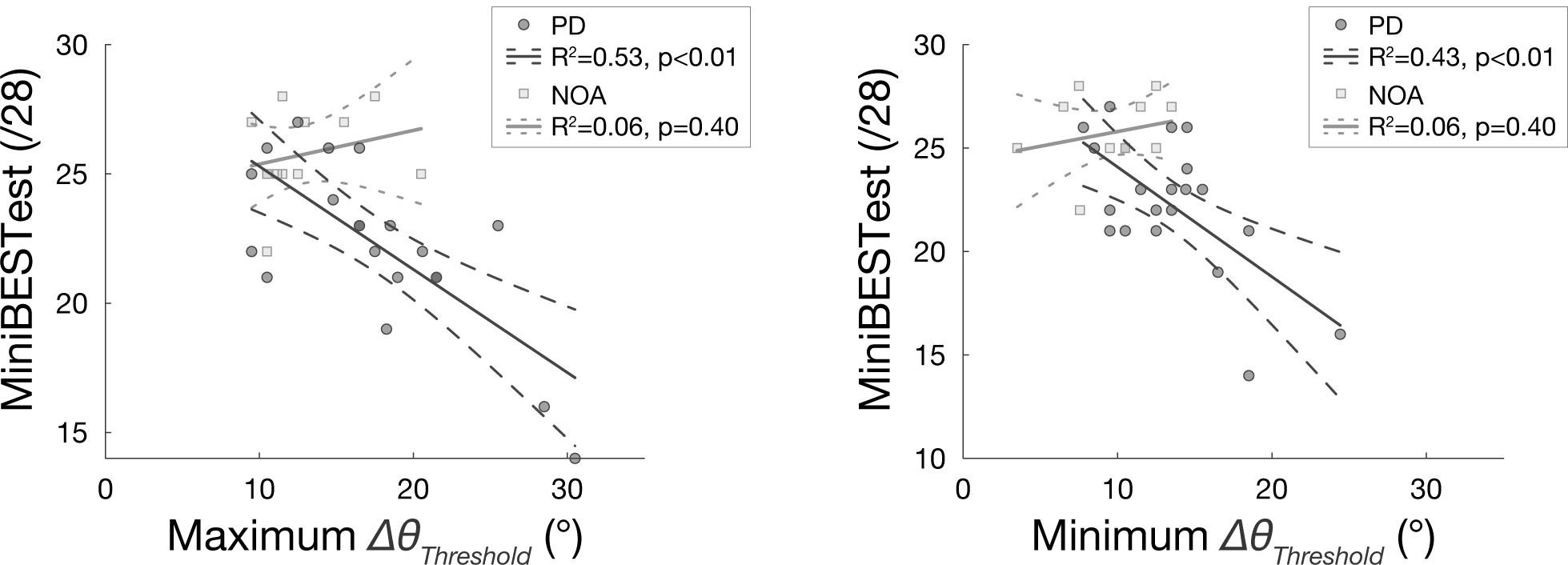
Associations between balance ability, as measured by MiniBESTest, and whole-body motion direction perception in NOA (squares) and PD (circles). Left: MaximumΔ*θ* _*Threshold*_. Right: Minimum Δ*θ* _*Threshold*_.

### Impaired directional acuity in people with PD is associated with symptom severity

Additional linear mixed models were implemented to more precisely determine associations between whole-body motion perception and PD severity, as measured either by MDS-UPDRS III score or by disease duration. Among PD participants, impaired perception of whole-body motion was associated with more severe MDS UPDRS III scores (maximum Δ*θ* _*Threshold*_: p=0.002; minimum Δ*θ*_*Threshold*_: p=0.063; Figure 4A). Positive, but statistically-insignificant associations were observed between threshold values and disease duration (maximum Δ*θ* _*Threshold*_: p=0.271; minimum Δ*θ* _*Threshold*_: p=0.770; Figure 4B). No significant differences were identified between thresholds on the more or less affected sides (p=0.74; Supplementary Materials S1).

**Figure 4.**
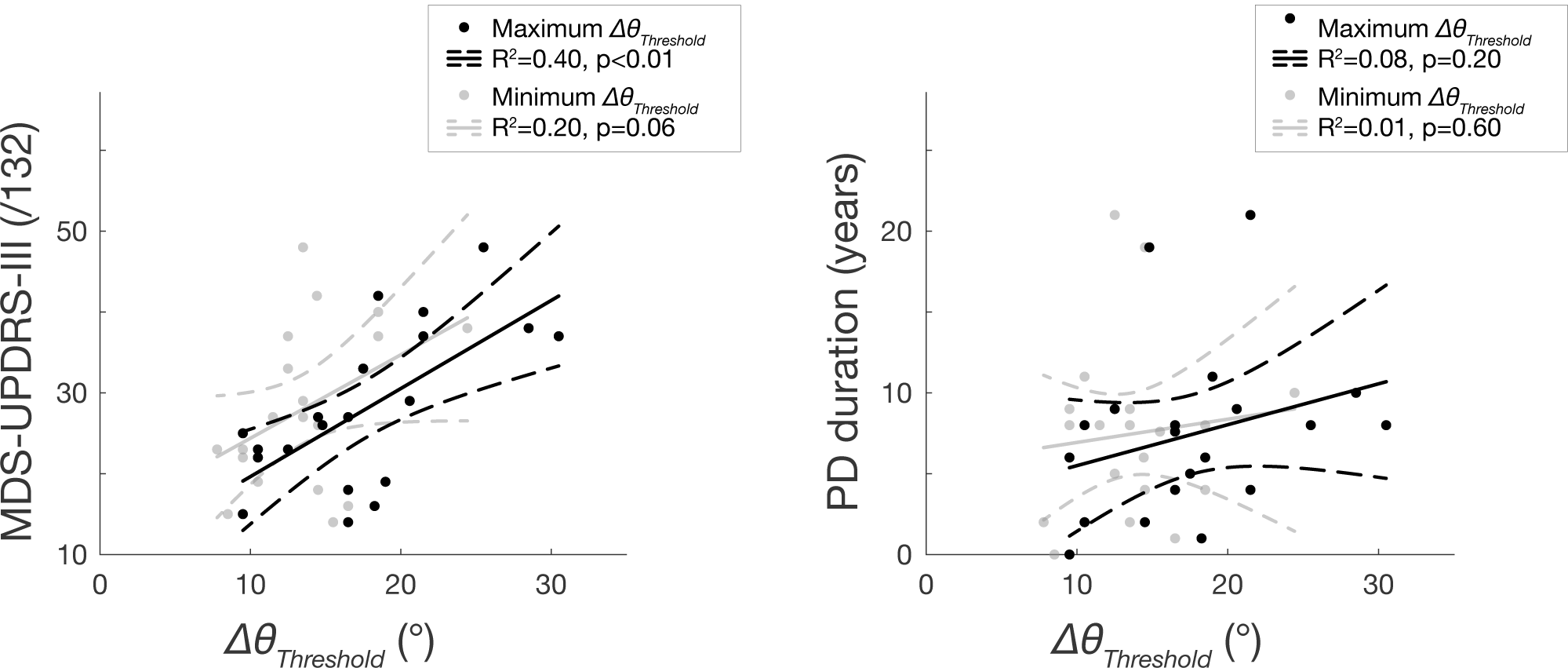
Associations between PD disease severity and disease duration and perception threshold among the PD group. Left: Disease severity. Right: Disease duration.

## Discussion

Here we identified a new functional perceptual deficit that may lead to falls in PD. To our knowledge, this is the first study investigating whole-body directional perception during standing in people with PD. Consistent with prior work demonstrating altered somatosensation, nociception, thermal sensation, and proprioception in PD [40], we show that directional acuity to whole-body motion imposed by support surface translations is also impaired. This perceptual deficit is significantly correlated with overall parkinsonian symptom severity, as determined by OFF medication MDS-UPDRS-III score, providing evidence that it may be associated with dopamine loss. This perceptual deficit is also significantly correlated with impaired balance among participants with PD, but not among NOA, supporting the hypothesis that it could represent an underlying cause of impaired balance in this population.

Taken together with data from an earlier report in young healthy individuals, these data show that directional perception of whole-body motion is affected by PD, but probably does not appreciably decline over healthy aging. We found no effect of age on directional perception in our current sample, which was limited to ages 56–82. We also found no significant differences between the current healthy sample and existing data of a young healthy sample (Supplementary Materials S1). However, we did find that individuals of very advanced age were more likely to be unable to tolerate the testing protocol, whether or not they had Parkinson’s disease. It is therefore possible that directional perception could decline rapidly at very advanced age, interfering with the testing protocol.

We were somewhat surprised to find no difference in perceptual acuity between the more- and less-affected side of participants with PD. We found a highly-significant association between discrimination threshold and parkinsonian symptom severity (MDS-UPDRS III score) (Figure 4), and a comparatively smaller effect of disease duration. Previously, grating discrimination threshold has been shown to be worse on the more affected side of people with PD [41], but the absence of a strong lateralization effect in this study could reflect the fact that most of these patients were ⩾ Hoehn & Yahr stage 2, with evident bilateral symptoms [42, 43]. It is also possible that this deficit may relate to axial symptoms, rather than to lateralized limb features. Consistent with somatosensory abnormalities that can occur early in disease progression [40], it is possible reduced whole-body direction acuity would be more lateralized in PD patients early in disease progression who tend to exhibit unilateral symptoms. However lateralization may be less evident in whole-body motion perception where directional acuity likely depends on sensory information from both sides of the body.

Our results suggest that estimates of whole-body directional acuity may be an important and specific contributor to falls in PD. We found an association between directional acuity and balance ability as measured by the MiniBEST test in PD and not NOA. However, in additional post hoc analyses, we found no statistically-significant associations between impaired directional acuity and fall history, which was available in a subset of participants (N=20; Supplementary Materials S1). However, maximum Δ*θ* _*Threshold*;_ had a ⩾30% stronger relationship to history of falls than the MiniBESTest [44]. We also found that patients who were unable to complete the testing procedure also had significantly reduced balance ability (p<0.01).

Our results are consistent with a growing body of evidence showing a variety of sensory deficits in PD that may underlie motor impairments. Prior studies of somatosensory deficits in PD have primarily examined isolated limbs or single joint tasks [2-8, 21, 24]. Here, we eliminated visual and auditory inputs, requiring participants to judge whole-body motion direction based only on somatosensory and vestibular inputs [45, 46]. The poor directional acuity in PD is consistent with less directionally-specific activation of automatic, brainstem-mediated balance-correcting muscle activity in PD [16, 17]. Additionally, reduced perceptual discrimination of whole body motion direction would also impair the efficacy of attentional mechanisms used to compensate for poor automatic balance control [47]. Similarly, deficits in body axial rotation perception during standing [24], and impairments in perceiving leg movement during walking [26] likely contribute to functional sensorimotor deficits.

Based on the strong association of whole-body perceptual deficits with MDS-UPDRS III score, we speculate that directional acuity would improve with effective symptomatic pharmacotherapy. In this study, all testing was performed ⩾12 hours after intake of dopaminergic medications. The perceptual deficit we identified here is strongly related to overall parkinsonian symptom severity, which generally respond well to acute dopamine challenge [48]. Several groups have also shown that impaired acuity to tactile stimuli improves with dopaminergic therapy [40]. Improving perceptual deficits that may cause falls and other motor deficit may be an additional benefit of careful management of “off”-times of dopaminergic fluctuations. Further PD-specific sensory deficits may contribute to the differences in parkinsonian fall circumstances compared to the general geriatric population [9].

## Acknowledgements

This study was supported in part by National Institutes of Health grants 2R01HD046922, R01 HD090642, K25HD086276, and the Sartain Lanier Family Foundation. The funders had no role in the conduct or publication of the study.

## Supporting Information

**Supplementary Materials S1.** Secondary analyses including Tables S1-S9 and Figure S1.

**Supplementary Data S1.** Source data used in statistical analyses.

