## Supplementary Materials S1 for "Perception of whole-body motion during balance perturbations is impaired in Parkinson’s disease and is associated with balance impairment"

### S1. Participant-level threshold values

Among threshold values included in analyses, 10/64 were determined using a psychometric curve fit to the PEST response data. Participant-level values are summarized in Table S1.

Table S1. Identified thresholds and characteristics of PD and NOA participants

| Code | Max Th (°) | Min Th (°) | Age | Sex | MiniBESTest (/28) | MDS UPDRS-III (/132) | MDS UPDRS-III Asymmetry Score | PD duration (years) | MoCA (/30) |
|---|---|---|---|---|---|---|---|---|---|
| PD01 | 30.5 | 18.5 | 56 | M | 14 | 37 | 0 | 8 | 27 |
| PD02 | 16.5 | 14.5 | 60 | M | 26 | 18 | -0.67 | 4.6 | 27 |
| PD03 | 17.5 | 12.5 | 59 | F | 22 | 33 | 0.68 | 5.5 | 27 |
| PD04 | 28.5 | 24.4‡ | 67 | F | 16 | 38 | 0.15 | 10 | 23 |
| PD05 | 18.3‡ | 16.5 | 56 | F | 19 | 16 | 0.2 | 1.2 | 29 |
| PD06 | 9.5 | 8.5 | 60 | F | 25 | 15 | -0.14 | 0.7 | 29 |
| PD07 | 16.5 | 11.5 | 62 | M | 23 | 27 | 0 | 7.8 | 28 |
| PD08 | 10.5 | 7.9‡ | 58 | M | 26 | 23 | 0.63 | 2.7 | 28 |
| PD09 | 19.0‡ | 10.5 | 65 | M | 21 | 19 | 0.17 | 11.9 | 23 |
| PD10 | 18.5 | 14.4‡ | 80 | M | 23 | 42 | 0.06 | 6.2 | 28 |
| PD11 | 21.5 | 12.5 | 64 | M | 21 | 37 | 0.08 | 21 | 25 |
| PD12 | 25.5 | 13.5 | 69 | M | 23 | 48 | -0.09 | 10 | 28 |
| PD13 | 14.5 | 13.5 | 69 | M | 26 | 27 | 0.09 | 2.7 | 27 |
| PD14 | 20.6‡ | 13.5 | 71 | M | 22 | 29 | 0.05 | 9.2 | 30 |
| PD15 | 14.8‡ | 14.5 | 72 | F | 24 | 26 | -0.06 | 19.7 | 29 |
| PD16 | 9.5 | 9.5 | 67 | M | 22 | 25 | -0.18 | 8.1 | 26 |
| PD17 | 10.5 | 9.5 | 75 | F | 21 | 22 | 0.18 | 8.2 | 24 |
| PD18 | 21.5 | 18.5 | 61 | F | 21 | 40 | 0.04 | 4.9 | 30 |
| PD19 | 12.5 | 9.5 | 56 | M | 27 | 23 | -0.41 | 9.8 | 30 |
| PD20 | 16.5 | 15.5 | 61 | F | 23 | 14 | 0.2 | 7.6 | 28 |
| NOA01 | 11.1‡ | 10.5 | 68 | F | 25 | - | - | - | 28 |
| NOA02 | 20.5 | 12.5 | 68 | M | 25 | - | - | - | 28 |
| NOA03 | 15.5 | 13.5 | 59 | F | 27 | - | - | - | 30 |
| NOA04 | 10.5 | 3.5 | 61 | F | 25 | - | - | - | 29 |
| NOA05 | 17.5 | 12.5 | 62 | F | 28 | - | - | - | 26 |
| NOA06 | 10.5 | 7.6‡ | 58 | M | 22 | - | - | - | 22 |
| NOA07 | 11.5 | 7.5 | 71 | F | 28 | - | - | - | 30 |
| NOA08 | 13.0‡ | 11.5 | 59 | F | 27 | - | - | - | 27 |
| NOA09 | 12.5 | 10.5 | 81 | M | 25 | - | - | - | 28 |
| NOA10 | 9.5 | 6.5 | 57 | M | 27 | - | - | - | 26 |
| NOA11 | 11.5 | 10.5 | 73 | F | 25 | - | - | - | 28 |
| NOA12 | 10.5 | 9.5 | 56 | F | 25 | - | - | - | 26 |

Max Th, Min Th: Maximum, Minimum thresholds of whole-body motion direction perception.
‡Threshold value estimated from psychometric curve fit.

### S2. Associations between identified threshold values and MiniBESTest score

Numerical results of linear mixed models examining associations between MiniBESTest score and directional acuity are presented in Table S2. Significant associations were identified among PD ($p<0.01$) but not among NOA ($p>0.43$).

Table S2. Associations between MiniBESTest score and whole-body motion perception among PD and NOA groups.

| Predictor | Beta coefficient | 95% CI | P value |
|---|---|---|---|
| PD | | | |
| Maximum threshold (point/°) | -0.40† | -0.59, -0.22 | <0.001** |
| Minimum threshold (point/°) | -0.53† | -0.84, -0.23 | 0.002 |
| NOA | | | |
| Maximum threshold (point/°) | 0.13 | -0.22, 0.48 | 0.43 |
| Minimum threshold (point/°) | 0.14 | -0.26, 0.54 | 0.45 |

**P<0.01, significant effect of threshold value on MiniBESTest score, linear mixed models.
†P<0.05, significant difference between PD and NOA, linear mixed models.

### S3. Validation of psychometric curve fit estimation

In cases where the PEST algorithm did not converge during data collection, we estimated the threshold using a psychometric curve fit to the available response data (*FitPsycheCurveLogit.m*). Thresholds from the psychometric curve fit were determined at 75% correct responses. The width of the curve (Figure S1A,B) was calculated as the width between $\Delta\theta$ values corresponding to 75% and 25% correct responses.

Linear regression analyses between $\Delta\theta_{Threshold}$ values identified via psychometric curve fits to available PEST response data and identified by convergence during the PEST procedure showed very strong linear relationships. Therefore, in cases where one threshold failed to converge during the PEST paradigm, the threshold was estimated as a linear conversion of the psychometric curve fit threshold using regression coefficients identified separately for each group and side (Table S3).

Table S3. Linear associations between $\Delta\theta_{Threshold}$ values identified using the PEST procedure and estimated from a psychometric curve fit to available data.

| Stratum | Slope | Intercept | $R^2$ |
|---|---|---|---|
| PD | | | |
| Left side | 1.10 | 0.03 | 0.93 |
| Right side | 0.95 | 0.92 | 0.91 |
| NOA | | | |
| Left side | 1.00 | 0.20 | 0.72 |
| Right side | 0.87 | 1.90 | 0.92 |

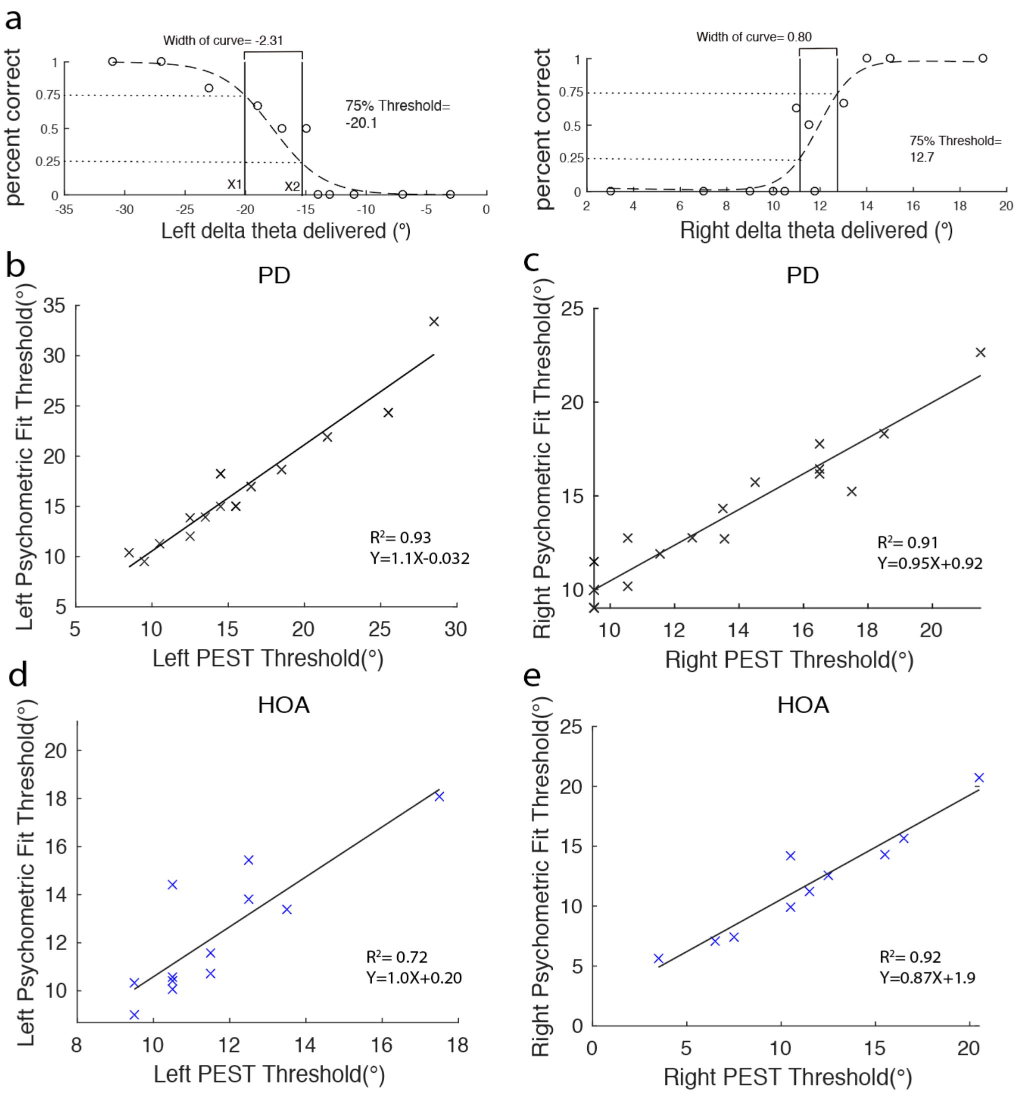


Figure S1. A. An example of psychometric curve fit of one of PD subject's PEST data, in which the proportion of correct trials are plotted vs. the tested deviation angle $\Delta\theta$. X1 and X2 denote deviation angles corresponding to 75% and 25% correct response rate, respectively. B-E. Thresholds determined by the PEST procedure were strongly linearly related to thresholds calculated from psychometric curve fit for b) left thresholds of PD subjects ($R^2$=0.93; y=1.1x-0.032), c) right thresholds of PD subjects ($R^2$=0.91; y= 0.95x+0.92), d) left thresholds of HOA subjects ($R^2$=0.72; y=1.0x+0.2), e) right thresholds of NOA subjects ($R^2$=0.92; y=0.87x+2).

### S4. Associations between MDS-UPDRS-III motor symptom asymmetry and threshold levels

We compared thresholds identified on the more- and less-affected side of PD participants to test whether asymmetries in whole-body motion directional acuity were associated with asymmetric symptoms. We classified each patient as left affected, right affected, or bilateral, respectively, as determined by an asymmetry score derived from the MDS-UPDRS III [38]:

$$Asymmetry\ Score = \frac{\sum MDSUPDRS_{iii}^{Left} - \sum MDSUPDRS_{iii}^{Right}}{\sum MDSUPDRS_{iii}^{Left} + \sum MDSUPDRS_{iii}^{Right}}$$

Positive values indicate that the more affected side is the left side, while zero indicates bilateral severity. N=18 and N=2 participants were categorized as asymmetric or bilateral, respectively.

We compared thresholds of the more and less affected sides of the N=18 asymmetric patients with a paired *t*-test. Identified thresholds did not differ across sides (more affected, 14.7±5.2°; less affected, 15.0±5.0°; p=0.74).

### S5. Associations between threshold magnitude and convergence

Thresholds estimated from psychometric curve fit on sides that converged were compared to thresholds estimated from psychometric curve fit on sides that did not converge during PEST experimental session within each group. In PD group, the mean and standard deviation of thresholds of converged sides was 14.4±4.1°, while the non-converged thresholds averaged at 10.4±5.3°. In NOA group, the mean and standard deviation of thresholds of converged and non-converged trials in HOA group were 11.5±4.8° and 10.4±2.7°, respectively. However, the difference between converged and non-converged thresholds within each group did not reach significance (HOA: p= 0.65, PD: p=0.29).

Additionally, the width of curve of psychometric curve fit in PD (1.52±1.85°) was greater than NOA (1.13±0.85°), where values further from 1.0 indicate worse discrimination sensitivity. However, the difference in the discrimination sensitivity between PD and NOA was not statistically-significant (p=0.33).

### S6. Comparison of directional acuity with an existing sample of young healthy adults

In order to assess associations between directional acuity and age, we compared identified threshold values $\Delta\theta_{Threshold}$ with existing data of healthy young adults (HYA; average age 22±3 y) collected previously [28]. Differences between PD, NOA, and HYA groups on threshold values were determined with separate one-way ANOVAs and Tukey post-hoc tests.

Average threshold values are summarized in Table S4. Compared to NOA, thresholds in HYA were similar overall but slightly larger in magnitude (0.4°, 3.5%). Significant main effects of group were identified for both Maximum Threshold (p=0.019) and Minimum Threshold (p=0.008). Post-hoc tests identified significant contrasts between PD and NOA (Minimum Threshold, p=0.040; Maximum Threshold, p=0.033) and between PD and HYA (Minimum Threshold, p=0.017; Maximum Threshold, p=0.070) but no differences between NOA and HYA (p≥0.96).

Table S4. Comparison of average threshold values in PD and NOA groups with existing data of young healthy participants (HYA).

| Variable | HYA N=11 | NOA N=12 | PD N=20 |
|---|---|---|---|
| Maximum (°) | | | |
| Mean±SD | 13.3±4.8 | 12.8±3.3[a] | 17.6±5.9[a] |
| Range | 8.5–23.5 | 9.5–20.5 | 9.5–20.5 |
| Minimum (°) | | | |
| Mean±SD | 10.0±3.4[b] | 9.7±2.9[c] | 13.5±4.0[b,c] |
| Range | 4.5–15.5 | 3.5–13.5 | 7.9–24.4 |

[a-c]Significant difference between marked groups, $P<0.05$, post-hoc tests.

### S7. Clinical and demographic variables associated with failure to complete testing protocol

We performed additional exploratory analyses post hoc to identify candidate clinical and demographic variables associated with failure to complete whole-body motion perception testing. The number of participants who could not complete testing in each group are summarized in Table S5. Participants for whom testing results were unavailable due to equipment problems (N=1, PD; N=1, NOA) were excluded from these analyses.

Among 46 participants for whom whole-body motion testing was attempted, testing was terminated early in 12 (26%). Testing was terminated early due to: inability to tolerate sensory deprivation (42%), fatigue (33%), and inability to understand instructions (25%). The frequency of early termination did not vary across groups (P=0.50, Fisher's exact test). The reasons for early termination did not vary across groups (P=0.28, Fisher's exact test).

Table S5. Summary of participants who completed and who did not complete the planned testing protocol.

| Outcome | PD N=30 | NOA N=16 | Total N=46 |
|---|---|---|---|
| Completed testing protocol | 20 (67) | 12 (75) | 32 (70) |
| Both thresholds identified | 13 (65) | 9 (75) | 22 (69) |
| One threshold identified | 7 (35) | 3 (25) | 10 (31) |
| Did not complete testing protocol | 9 (30) | 3 (19) | 12 (26) |
| Could not tolerate sensory deprivation | 4 (44) | 1 (33) | 5 (42) |
| Fatigue | 4 (44) | 0 (0) | 4 (33) |
| Could not understand instructions | 1 (12) | 2 (66) | 3 (25) |
| Equipment failure | 1 (3) | 1 (6) | 2 (4) |

Frequencies are presented as N (%).

Separate logistic regression analyses were performed to identify candidate clinical and demographic variables associated with early testing termination among the 46 participants for whom testing was attempted. Continuous variables were transformed to *z*-scores prior to entry in logistic regression models. Associations between candidate predictor variables and inability to complete testing were expressed as odds ratios and confidence intervals (OR ± 95% CI; Table S6. These models identified significant effects of age (P=0.019) and impaired performance on

MiniBESTest (P=0.017) but not of presence of PD (P=0.439) or other variables related to PD severity on early termination of testing.

Table S6. Associations between demographic and clinical features and early termination of whole-body motion testing protocol.

| Predictor | OR | 95% CI | P value |
|---|---|---|---|
| Increased age | 2.55 | 1.17–5.56 | 0.019* |
| Female sex | 0.69 | 0.50–8.00 | 0.327 |
| Poorer MiniBESTest score | 2.46 | 1.17–5.15 | 0.017* |
| Presence of PD | 1.80 | 0.41–7.98 | 0.439 |
| Increased PD duration | 1.60 | 0.73–3.54 | 0.244 |
| Increased MDS-UPDRS-III total score | 1.57 | 0.70–3.65 | 0.290 |
| Increased MDS-UPDRS-III asymmetry | 0.10 | 0.00–2.17 | 0.142 |

*P<0.05.

### S8. Clinical and demographic variables associated with partial convergence during testing

We also performed exploratory analyses to identify candidate clinical and demographic variables associated with incomplete convergence on one side during the planned testing protocol. The number of patients who completed the planned testing protocol and for whom one or both thresholds were identified are summarized in *Section S7. Clinical and demographic variables associated with failure to complete testing protocol*.

Among 32 participants for whom the planned testing protocol was completed, estimates of one threshold failed to converge during the planned testing period in 10 (31%). These thresholds were subsequently estimated using a psychometric curve fit to the PEST results. The frequency of identifying only one threshold did not vary across PD and NOA (P=0.71, Fisher's exact test).

Similar to analyses reported for failure to complete testing, we performed additional logistic regression analyses to identify clinical and demographic factors associated with incomplete convergence during the planned testing protocol. Associations between candidate predictor variables and incomplete convergence were expressed as odds ratios and confidence intervals (OR ± 95% CI; Table S7). Logistic regression models identified no significant associations between clinical and demographic variables and incomplete convergence.

Table S7. Associations between demographic and clinical features and incomplete convergence during threshold identification.

| Predictor | OR | 95% CI | P value |
|---|---|---|---|
| Increased age | 1.23 | 0.58–2.60 | 0.58 |
| Female sex | 1.00 | 0.22–4.46 | 1.00 |
| Poorer MiniBESTest score | 1.84 | 0.69–4.89 | 0.22 |
| Presence of PD | 1.62 | 0.33–7.80 | 0.56 |
| Increased PD duration | 1.27 | 0.46–3.49 | 0.65 |
| Increased MDS-UPDRS-III total score | 0.92 | 0.28–3.07 | 0.90 |
| Increased MDS-UPDRS-III asymmetry | 0.87 | 0.27–2.78 | 0.81 |

### S9. Associations between identified threshold values and fall history

Additional analyses were performed to determine associations between identified threshold values and fall history among the PD sample. The number of falls over 6 months prior to study enrollment were recorded during interview, and participants were coded as "those with ⩽1 fall" and "recurrent fallers," as previously [30]. Summary statistics and standardized odds ratios and confidence intervals (OR ± 95% CI) for each predictor variable are presented in Tables S8 and S9, respectively. Logistic regressions identified no statistically-significant associations between Maximum Threshold, Minimum Threshold, or MiniBESTest score and faller status. However, positive associations with fall history were identified for each continuous variable examined, with the strongest association identified for Maximum Threshold.

Table S8. Comparison of average threshold values and MiniBESTest scores between PD participants with and without fall history.

| Number of falls in preceding 6 months | N | Max Th (°) | Min Th (°) | MiniBESTest (/28) |
|---|---|---|---|---|
| Any | 20 | 17.6±5.9 | 13.5±4.0 | 22.3±3.3 |
| 0 or 1 | 13 (65) | 16.1±5.5 | 13.3±4.5 | 22.8±2.8 |
| 2 or more | 7 (35) | 20.5±6.0 | 13.8±3.3 | 21.1±4.0 |

Table S9. Associations between identified threshold values, MiniBESTest scores, and fall history. Each predictor variable was transformed to a *z*-score prior to analysis.

| Predictor | OR | 95% CI | P value |
|---|---|---|---|
| Max Th (°) | 2.34 | 0.79–6.95 | 0.125 |
| Min Th (°) | 1.14 | 0.45–2.90 | 0.785 |
| MiniBESTest (score) | 1.76 | 0.64–4.84 | 0.273 |
